# Protein engineering approach to enhance activity assays of mono-ADP-ribosyltransferases through proximity

**DOI:** 10.1101/2022.04.11.487883

**Authors:** Albert Galera-Prat, Juho Alaviuhkola, Heli I. Alanen, Lari Lehtiö

## Abstract

Human mono-ADP-ribosylating PARP enzymes have been linked to several clinically relevant processes and many of these PARPs have been suggested as potential drug targets. Despite recent advances in the field, efforts to discover such compounds have been hindered by the lack of tools to rapidly screen for high potency compounds and profile them against the different PARP enzymes of the ARTD family. We here expanded the methods and engineered mono-ART catalytic fragments to be incorporated into a cellulosome-based octavalent scaffold. Compared to the free enzymes, the scaffold-based system results in an improved activity for the tested PARPs due to improved solubility, stability and the proximity of the catalytic domains, altogether boosting their activity beyond 10-fold in the case of PARP12. This allows us to measure their enhanced activity using a simple and easily accessible homogeneous NAD^+^ conversion assay, facilitating its automation to reduce the assay volume and lowering the assay costs. The approach will enable the discovery of more potent compounds due to increased assay sensitivity and it can be applied to compound screening campaigns as well as inhibitor profiling.

## Introduction

ADP-ribosylation is a phylogenetically ancient post-translational modification that participates in a wide range of key cellular processes (Perina *et al*., 2014). This modification is carried out by ADP-ribosyltransferases (ARTs) that transfer ADP-ribose from NAD^+^ to macromolecules. In humans, the largest group of ARTs is composed by 17 proteins of the ARTD family that share a conserved catalytic domain with homology to the diphtheria toxin-like bacterial ARTs (Lüscher *et al*., 2021). ARTDs can be classified in two groups according to the post-translational modification that they catalyse: mono-ARTs (PARP3-4, 6-12 and 14-16) transfer a single ADP-ribose unit to their substrates, while poly-ARTs (PARP1-2, TNKS1-2) are capable of sequentially transferring multiple ADP-ribose units to produce a linear or a branched polymer. The mono-ART group actually comprises most of the PARPs. Nevertheless, compared to poly-ARTs, they have been less studied and have only recently been the focus of drug discovery efforts (Challa *et al*., 2021).

PARP10 was the first identified human mono-ART (Yu et al., 2005; Kleine et al., 2008). Since then, several studies have reported multitude of functions for the different mono-ARTs including cancer biology (Vyas and Chang, 2014; Schleicher *et al*., 2018), viral replication (Atasheva *et al*., 2012; Verheugd *et al*., 2013), autophagy (Kleine *et al*., 2012; Carter-O’Connell *et al*., 2016), apoptosis (Herzog *et al*., 2013; Saei *et al*., 2021), inflammation (Welsby *et al*., 2014), immunology, transcription, splicing, translation, RNA degradation (Bock *et al*., 2015), and regulation of exocytosis (Grimaldi *et al*., 2020). Many mono-ARTs have been suggested as potential drug targets (Scarpa *et al*., 2013; Nicolae *et al*., 2014; Challa *et al*., 2021; Hopp and Hottiger, 2021) but the wide repertoire of functions of the different enzymes also stresses the importance of developing selective inhibitors that target specific enzymes. Such compounds could also be used in basic research to investigate in detail the function of these enzymes. Nevertheless, efforts to discover such compounds have been hindered by the lack of tools to rapidly screen for high potency compounds and profile them against the different enzymes (Glumoff *et al*., 2022).

Several assays to study ARTs activity based on biophysical or biochemical methods and employing either natural, radioactive or modified NAD^+^ (for a comprehensive review see (Glumoff *et al*., 2022)). The requirements for an ideal assay for drug discovery are its biological relevance, reproducibility, and quality. Furthermore, since screening is performed in singlets, signal robustness is also a key factor, typically determined according to the Z’ factor (Zhang *et al*., 1999; Hughes *et al*., 2011). Additionally, the assay should not be sensitive to the effects of the solvent or other components used. Finally, the use of small assay volumes and inexpensive reagents should be favored to reduce the costs, particularly when screening large compound libraries (Hughes *et al*., 2011).

Putt and Hergenrother developed a homogeneous assay based on quantifying the leftover NAD^+^ after the incubation with poly-ART PARP1 (Putt and Hergenrother, 2004), that was later adapted for its use with tankyrases as well as mono-ARTs (Narwal *et al*., 2012; Venkannagari *et al*., 2013) and it has been shown to be suitable for inhibitor screening. Upon reaction with potassium hydroxide, acetophenone and formic acid, NAD^+^ is converted to a fluorophore that can be quantified in a plate reader with excitation and emission wavelengths of 372 nm and 444 nm, respectively. This assay is generally robust, very cost-effective, homogeneous, and uses the natural substrate. Still, in the case of mono-ARTs, its biggest drawback is that it requires high enzyme concentration to reach sufficient NAD^+^ conversion. This does not only result in higher protein consumption but also limits the ability to resolve potent inhibitors as the lowest measurable IC_50_ value is equal to half of the protein concentration used.

Wigle and colleagues recently reported a new high-throughput assay for screening and profiling of mono-ART inhibitors (Wigle *et al*., 2020). This method is based on immobilizing the enzymes on microplates prior to their incubation in the presence of biotin-NAD^+^, which allows to quantify their level of automodification using dissociation-enhanced lanthanide fluorescence immunoassay (DELFIA). The assay is applicable to all the human mono-ARTs and allowed to screen and determine IC_50_ values, even for potent compounds. Nevertheless, the relatively high cost of the Ni^2+^-NTA-coated plates and the reagents required for the assay may make it in many laboratories a less suitable option for screening large compound libraries. Interestingly, although the performance of the assay can be partially attributed to the highly sensitive detection method, the proximity of the catalytic domains induced by their immobilization allowed to overcome the high K_M_ for self-modification and significantly boosted the enzymatic activity of the mono-ARTs.

Based on this observation we sought to improve the homogeneous NAD^+^ consumption assay method that is simple and easily accessible by promoting the proximity of the enzymes using a protein engineering strategy. Several methods to induce enzyme proximity at a protein level have been described (Dong *et al*., 2021) including fusion to multimeric protein (Mitsuzawa *et al*., 2009) or incorporation on scaffolds mimicking naturally occurring multi-protein complexes (Gad and Ayakar, 2021), such as the cellulosome complex used in this work. Cellulosomes are multi-enzyme complexes produced by several anaerobic bacteria to degrade plant cell-wall biomass (Bayer *et al*., 2004). These self-assemble on scaffolds composed of cohesin modules that interact with dockerin domain containing enzymes (Bayer et al., 2004). The modularity of these components, together with their high affinity (Slutzki et al., 2012) and existence of different specificity groups (Haimovitz et al., 2008) make them very suitable to generate well-defined multi-protein complexes generally known as designer cellulosomes (Bayer et al., 1994; Fierobe et al., 2001). This technology has been shown to be an invaluable tool to study cellulosome activity and the principle has also been applied to other fields such as biosensor development (Hyeon et al., 2014), as well as to improve of reactions taking advantage of the proximity induced by assembly on the scaffold (You et al., 2012; Liu et al., 2013; You and Zhang, 2014).

In this work we convert different mono-ARTs catalytic fragments into the cellulosomal mode for their incorporation into an octavalent designer scaffold in order to induce proximity (**Fig. 1**). This allows us to measure their enhanced activity using the homogeneous NAD^+^ conversion assay, which provides benefits in avoiding washing steps and the use of an inexpensive natural NAD^+^ as a substrate. The development started with PARP10 for which we have recently discovered compounds more potent than what the standard homogeneous assay allows us to measure (reported elsewhere). We describe here first the set up and validation for the PARP10 assay and then extend this approach to other human mono-ARTs.

**Fig. 1:**
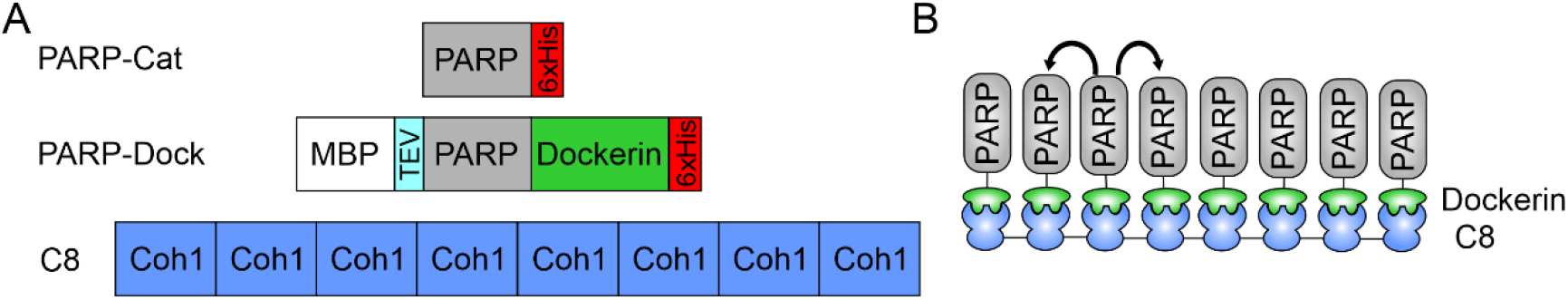
Construct design for the cellulosome-like scaffold-based assay. **(A)** Schematic representation of PARP-Cat, PARP-Dock constructs and C8 scaffold used in this work. **(B)** The C8 construct consisting of 8 tandem copies of cohesin domain provide the scaffold where dockerin carrying enzymes can be bound and brought to close proximity to facilitate modification of nearby units.

## Materials and methods

### Cloning

Dockerin containing proteins were cloned as follows: First, *Hungateiclostridium thermocellum* Cel48S dockerin, human SRPK2 and the catalytic fragment of human PARP10, PARP12, PARP16 were PCR amplified separately. The fragments containing SRPK2 or catalytic fragments were then PCR fused to dockerin and cloned using SLIC to pNIC-MBP (Sowa *et al*., 2020). Due to dockerin degradation in these constructs, the region containing MBP-PARP/SRPK2-Dockerin was PCR amplified and cloned into pNIC-CH using SLIC. Constructs were verified by sequencing.

MBP-SRPK2-dockerin plasmid was used as a template to generate the remaining clones. Briefly, MBP-SRPK2-Dockerin plasmid was linearized between Dockerin and MBP to allow SLIC cloning of PCR amplified catalytic fragments of PARP6, PARP7, PARP11, PARP14 and PARP15. Scaffolds C1 and C8 were described in a previous work (Galera-Prat *et al*., 2018). Details of the constructs can be found in **Table S1**.

### Protein expression and purification

The specific conditions for production and purification of proteins are summarized in **Table S2**. All proteins were expressed in *E. coli* BL21 (DE3), Rosetta 2 (DE3) or C41 (DE3) cells. Cells were grown in 1-3 L of Terrific Broth media supplemented with 50 μg/ml kanamycin (and 34 μg/ml chloramphenicol in the case of Rosetta 2 cells) and 0.8% glycerol to an OD600 of 1 - 1.2 before transferring the culture to expression conditions (**Table S2**). For dockerin containing constructs the media was supplemented with 2 mM CaCl_2_ at the beginning of the expression step. Cells were harvested by centrifugation at 5020 x g and resuspended in lysis buffer containing 30 mM HEPES pH 7.5, 500 mM NaCl, 10 mM Imidazole, 10% glycerol and 0.5 mM TCEP and supplemented with Pefabloc SC (Roche) before freezing in liquid nitrogen. Lysis of the cells was performed by addition of Lysozyme and DNAse I (Sigma-Aldrich) followed by sonication (Branson 250 Sonifier) for 2 minutes (5 s on, 15 s off) and centrifugation at 16000xg to remove cell debris. Clarified samples were then purified by Immobilized Metal Affinity Chromatography (IMAC) where the proteins were bound to the IMAC resin pre-equilibrated in lysis buffer, washed in a similar buffer containing 25 mM imidazole and eluted using 200-350 mM imidazole. Samples were then further purified with size exclusion chromatography (SEC) using S200 HiLoad sepharose 16/60 column (Cytiva) in SEC buffer containing 30 mM HEPES, pH 7.5, 350 mM NaCl, 10% glycerol, 0.5 mM TCEP. For Dockerin containing proteins, all buffers used during purification were supplemented with 2 mM CaCl_2_. Fractions were analysed by SDS-PAGE and selected fractions concentrated using Amicon15 (Millipore) to small volume before aliquoting and freezing. Protein concentrations were estimated using absorbance at 280 nm and the calculated extinction coefficient of the protein. An additional purification step using MBP-Trap (Cytiva) between the IMAC and SEC steps was performed for PARP6-Dock. Binding and washes to MBP-Trap column was performed in SEC buffer, while elution was done in the same buffer supplemented with 10 mM maltose.

Scaffolds were purified as previously described (Galera-Prat *et al*., 2018). Briefly, clarified samples were transferred to a new tube and they were incubated at 55°C for 30 min followed by 10 min in ice. The heat-treated samples were then subjected to centrifugation at 4000xg for 10 min, the supernatants were filtered through 0.45 μm filter and purified using 5 ml IMAC column (Cytiva) as described above. The eluted sample was buffer exchanged to SEC buffer using Amicon 15, concentrated and stored at −70°C. MBP protein was obtained from PARP10-Dock after overnight TEV cleavage and IMAC purification, followed by a SEC exclusion step.

Dockerin was purified as a fusion protein to xylanase T6 from *Geobacillus stearothermophilus* containing a TEV protease site between xylanase and dockerin domains as well as a His tag at the C-terminus described elsewhere (Vera et al., 2021). After IMAC and SEC, the protein was incubated with 1:30 TEV protease overnight and the resulting sample was further purified using IMAC column to recover the cleaved dockerin fragment.

### Analytical size exclusion chromatography

S200 increase (Cytiva) was used for analytical size exclusion chromatography. 35 μM of enzyme was mixed with different ratios of scaffold in SEC buffer with 2 mM CaCl_2_ and incubated in ice for 1 h before injection. Fractions were collected and further analysed by SDS-PAGE.

### Determination of complex molecular weight

Enzyme (35 μM) was mixed with scaffold in SEC buffer with 2 mM CaCl_2_. The sample was incubated in ice for 1 h before injection to S200 Increase 10/300 (Cytiva) column connected to a Minidawn multiangle light scattering (MALS) and RI detectors (Wyatt). Concentration was measured from RI detector and data was analyzed with Astra 7.3.2 (Wyatt Technology).

### ADP-ribosylation assay and inhibitor concentration response measurements

Concentrations refer to that of the catalytic unit. Unless otherwise stated, the concentration of C8 scaffold used in the assays was selected based on the experimentally determined optimal ratio to the enzyme used, while C1 scaffold was added at the same molar concentration as the catalytic unit. When SRPK2 was used in an assay, it was also included in the NAD^+^ control in order to correct for its effect to the fluorescence.

The different components were mixed manually or in a semi-automated manner using Echo (Beckman Coulter) for small compounds and Mantis (Formulatrix) on 96 or 384 well plates. Plates were incubated for specified times at 300 rpm and room temperature on Biosan plate shakers. The reaction was stopped by addition of 2 M KOH before addition of 20% acetophenone for 10 min and formic acid. The resulting fluorescence was measured using Infinite M1000 or Spark plate readers (Tecan). Initial analysis of the data was performed in excel, while plotting and fitting was performed in GraphPad Prism version 9.3.0. Concentration response reactions were carried out in quadruplicates and sigmoidal IC_50_ curves were fitted using four variables.

### Western blot

5 μg of PARP (6-7 μM depending on the particular enzyme) in 10 μl volume was incubated overnight at room temperature in the presence of 50 mM NAD^+^, and 10 μMSRPk2, 0.7-0.8 μM C8, 6-7 μM C1, 6-7 μM MBP or 6-7 μM Dockerin. After incubation, 2 μl of a 1:10 sample dilution was loaded to SDS-PAGE and analysed by western blot with nLuc-eAF1521 as detection method as described elsewhere (Sowa *et al*., 2021). Briefly, samples were separated by SDS-PAGE, then transferred to nitrocellulose membrane using semi-dry trans blot system (Biorad). The resulting membranes were first stained with Ponceau red (Biorad), de-stained in TBS + 0.1% tween20 and blocked in 5% milk. Membranes were incubated in TBS + 0.1% Tween 20 + 1 % milk containing 1 μg/ml of nLuc-eAF1521 for 15 min at 4°C. 5 min washes were repeated 3 times with TBS-T before addition of nano-Glo diluted 1:1000 in 10 mM sodium phosphate pH 7.5 solution for development. Imaging of the membrane was performed in a GelDoc (Biorad) system.

## Results

### System design and planning

Incorporation to a cellulosome-like scaffold composed of cohesin module repeats can be achieved by fusing a dockerin domain to the different PARP catalytic domains. For this work we used an artificial scaffold composed of 8 repeats of *H. thermocellum* CipA cohesin 1 (Galera-Prat *et al*., 2018). Cel48S is the major exocellulase from *H. thermocellum* cellulosome and its dockerin was selected due to its capacity to bind CipA cohesin 1 (Lytle *et al*., 1996). Fusion proteins were designed to contain Cel48S dockerin fused at the C-terminus of the different PARP catalytic domains. Previous PARP10 constructs contained a C-terminal His tag, indicating that it is possible to fuse additional sequence to that part of the protein and additionally it results in a construct were dockerin is placed in its natural location. Additionally, all dockerin-containing constructs present a C-terminal tag that facilitates their purification and a TEV-cleavable N-terminal MBP tag to improve solubility and to provide additional purification options (**Fig. 1A**).

### Complex formation and scaffold ratio determination

We first determined the complex formation capacity of the purified proteins. For this, we analysed PARP10-Dock in SEC to experimentally determine the concentration at which all available PARP10-Dock was incorporated into the complex (**Fig. 2A**). Due to the low extinction coefficient of the scaffold and presence of impurities in the sample (**Fig. S1**), the concentration determination was not very accurate which hampered accurate determination of the exact stoichiometry of the complex. To overcome this, we analysed some of the complexes with SEC-MALS for direct determination of their molecular weights. Molecular weight of the complexes determined at the ratio that resulted in no visible unbound PARP10-Dock was found to be 650 ± 4 kDa (**Fig. 2B**), that closely matches the expected molecular weight of a C8 scaffold loaded with 7 PARP10-Dock (656 kDa). Apparent excess of PARP10-Doc resulted in a complex of 714 ± 9 kDa, closer to the fully occupied complex with expected molecular weight of 731 kDa (**Fig. 2C**). Taken together this data indicate that the scaffold is able to bind up to 8 PARP10-Dock at a time, although the preferred complex contains 7 enzymatic units.

**Fig. 2.**
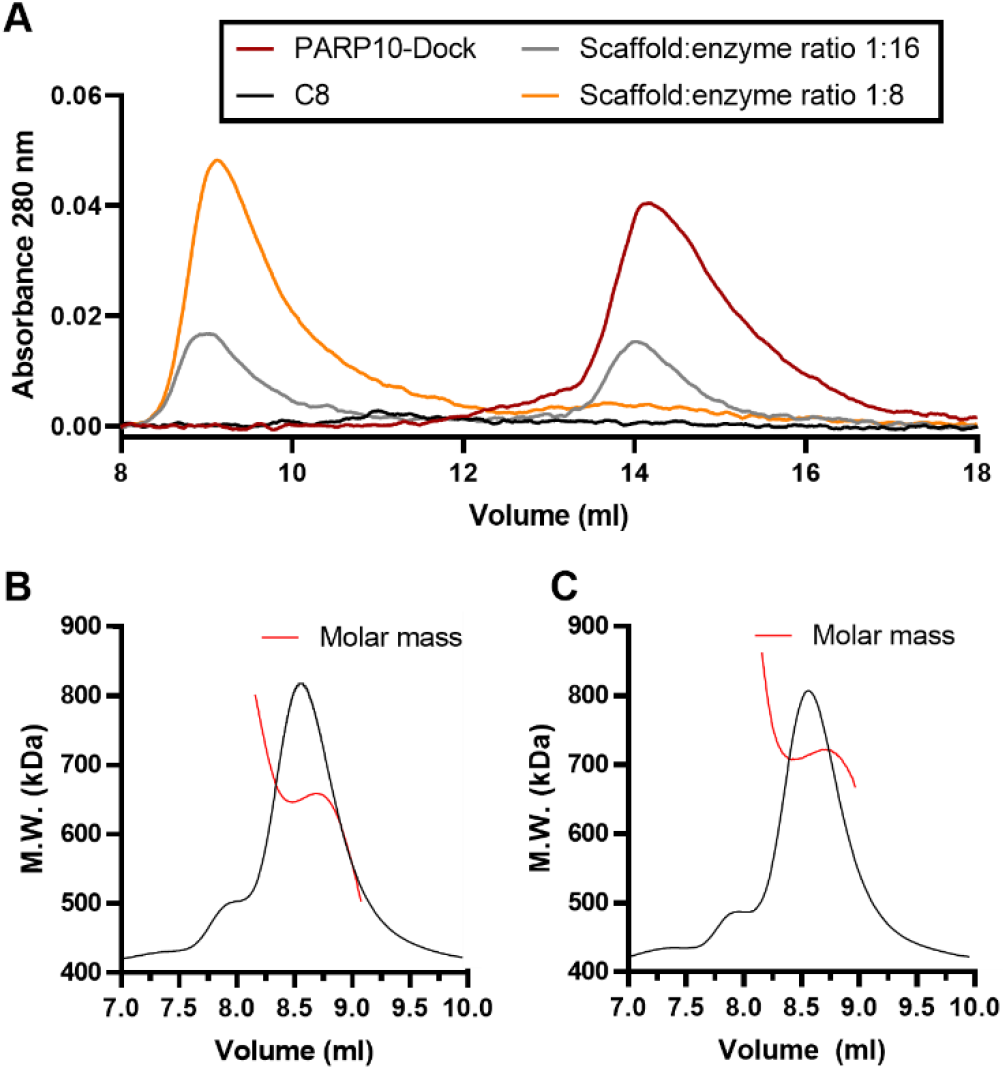
Control experiments for the assembly. **(A)** SEC profile of PARP10-Dock mixed with different ratios of C8 scaffold. **(B)** SEC-MALS analysis of PARP10-Dock and C8 scaffold under saturating conditions. The determined molecular weight of the complex is 650 ± 4 kDa. **(C)** SEC-MALS analysis of PARP10-Dock with limited C8 scaffold showing a molecular weight of 714 ± 9 kDa.

### Enhanced solubility and proximity contribute to improved PARP10-Dock activity

First we studied the NAD^+^ conversion capacity of PARP10-Dock upon titration of the scaffold to determine the optimal enzyme:scaffold concentration (**Fig. 3A**). The resulting curve shows a clear maximum at the same ratio where SEC confirmed the incorporation of all enzymatic units to the scaffold. This ratio was used for all subsequent experiments and to correct the scaffold concentration for all the assays.

**Fig. 3.**
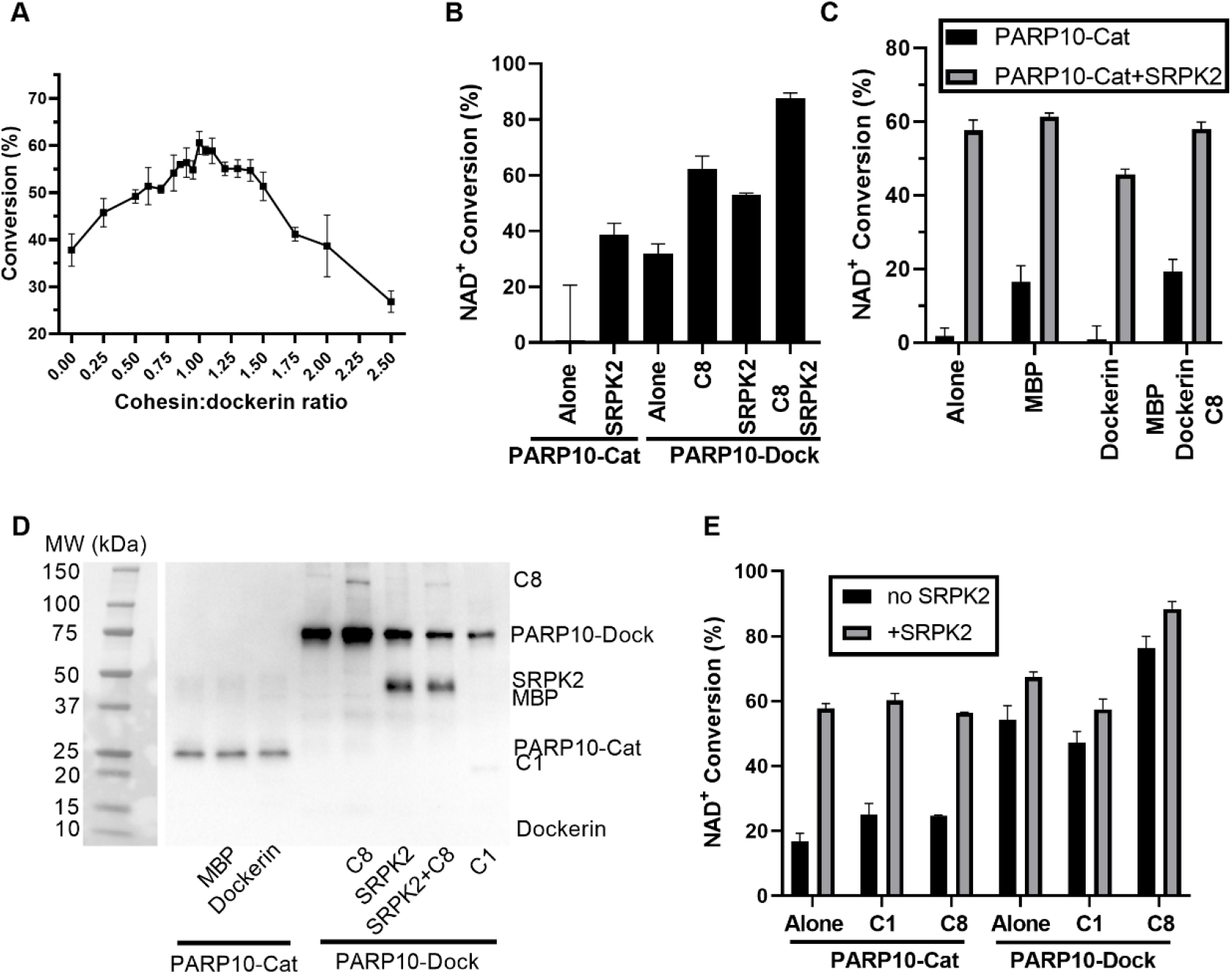
NAD^+^ conversion activity of PARP10-Dock incorporated to the scaffold. **(A)** PARP10-Dock NAD^+^ conversion activity as a function of scaffold ratio. Experiment performed with 100 nM PARP10-Dock in the presence of 500 nM SRPK2 and incubated for 16 h. **(B)** Comparison of the original and scaffold based homogeneous assay for PARP10 at 200nM enzyme and 2μM SRPK2 for 4h. **(C)** Effect of MBP and dockerin on the NAD^+^ conversion of PARP10-Cat, performed at 300 nM enzyme and 2 μM SRPK2 for 4h. **(D)** Western blot analysis of MARylated components during the assay. **(E)** The proximity induced by the C8 scaffold increases the NAD^+^ conversion of the system measured with 100 nM enzyme, equimolar cohesin:dockerin concentrations and 2 μM SRPK2 for 4h. NAD^+^ conversion values correspond to average ± SD of 4 repeats.

NAD^+^ consumption assay revealed that PARP10-Dock presents higher and more consistent NAD^+^ conversion capacity than PARP10-Cat (32 ± 3% and 0.8 ± 20%, respectively) and comparable to that of the original assay with PARP10-Cat in the presence of SRPK2, a target protein for ADP-ribosylation (39 ± 4%, **Fig. 3B**). Incubation of PARP10-Cat with free MBP or dockerin, the additional elements present in PARP10-Dock, only showed increased activity with MBP (**Fig. 3C**). Under these conditions, the NAD^+^ conversion was still lower compared to that of PARP10-Dock and the effect was negligible when SRPK2 was present. Western blot using nanoLuc-eAF1521 as detecting agent showed no modification of MBP or dockerin by PARP10-Cat (**Fig. 3D**). Altogether, these results suggest that neither MBP nor dockerin serve as additional MARylation targets, but that the presence of MBP can promote PARP10-Dock solubility or stability resulting in enhanced activity.

The presence of C8 scaffold resulted in a clear increase in NAD^+^ conversion activity of PARP10-Dock (54 ± 4% vs. 76 ± 3% upon scaffold addition), while scaffold C1 led to a small decrease (47 ± 3%, **Fig. 3E**). Since the only difference between C1 and C8 scaffolds is that the latter can induce the formation of complexes were multiple PARP10-Dock bind to the same scaffold polypeptide, it is clear that the scaffold induced proximity can improve PARP10 NAD^+^ conversion. As expected, addition of either scaffold to PARP10-Cat did not affect its NAD^+^ conversion activity (**Fig. 3E**). Western Blot analysis of PARP10 self-modification in the presence of scaffold shows a similar trend, where the presence of C8 results in enhanced auto-modification of PARP10-Dock compared to PARP10-Cat, while it is reduced in the presence of C1 and does not affect PARP10-Cat (**Fig. 3D**). This analysis also revealed that the scaffolds can be modified both by PARP10-Cat and PARP10-Dock, although the level of modification is low. This is in agreement with the negligible change in NAD^+^ conversion observed for PARP10-Cat in presence of C1 and C8 (**Fig. 3E**), the contribution of this to the overall NAD^+^ conversion appears to be negligible. All together, these results indicate that the formation of large complexes where multiple PARP10 catalytic domains are brought together can increase the overall conversion of NAD^+^ in a homogeneous assay by increasing the self-modification of PARP10.

SRPK2 is a natural PARP10 substrate and its incorporation in the homogeneous NAD^+^ conversion assay resulted in increased PARP10 NAD^+^ conversion and therefore allows to reduce the needed PARP10 concentration (Venkannagari *et al*., 2016). We observed that SRPK2 induced an additional increase in NAD^+^ conversion under all conditions (**Fig. 3B, 3C, 3E**), even when PARP10-Dock was forming a complex with C8 scaffold. The enhanced NAD^+^ conversion of the scaffold-based assay can be exploited to lower the necessary PARP10 concentration and/or the duration of the assay.

### Optimization and validation of PARP10-Dock scaffold-based assay

We next sought to optimize the conditions of the assay. The target was to reduce the concentration of PARP10-Dock to as low as possible while keeping the assay robust, simple, short and that can be reliably prepared using automated dispensing robots, so that screening large compound libraries is feasible. Despite the use of SRPK2 can improve the activity of the system, dispensing it automatically has shown inconsistent results. Since in the absence of SRPK2 the scaffold-based system already provides an improved activity, we targeted at optimizing the assay in two forms: a low volume version without SRPK2 that can be dispensed automatically targeted for measuring IC_50_ of large number of compounds, and a very low protein concentration version with SRPK2 that can be used for profiling and studying high potency compounds.

We set a target of 30% total conversion to avoid conversion rate to be affected by substrate depletion (**Fig. 4A**) and 4 h assay to find useful PARP10-Dock concentration (**Fig. 4B**). This allowed to determine a working concentration of 150 nM of PARP10-Dock in 10 μl for the automatic assay (**Table I**). SRPK2 concentration showed a strong effect on the activity of the system (**Fig. 4C**) and in the presence of 3 μM SRPK2, the assay can be carried out with 20 nM PARP10-Dock in 50 μl in the manual assay (**Table I**). These conditions represent a clear performance increase of the assay compared to the original homogeneous assay were 50 μl of 150 nM PARP10-Cat had to be manually mixed and incubated for 16 h to achieve comparable conversion (Venkannagari *et al*., 2016). The use of both conditions was validated by measuring dose-response curves for OUL35 compound, yielding IC_50_ measurements of 417 nM in the low protein concentration manual assay (**Fig. 4D**; 95% CI 190 - 980 nM) and 414 nM for the automated mode (**Fig. 4E**; 95% CI 210 - 820 nM). Both values are well in agreement with the reported IC_50_ for PARP10-Cat of 330 nM (Venkannagari *et al*., 2016).

**Fig. 4.**
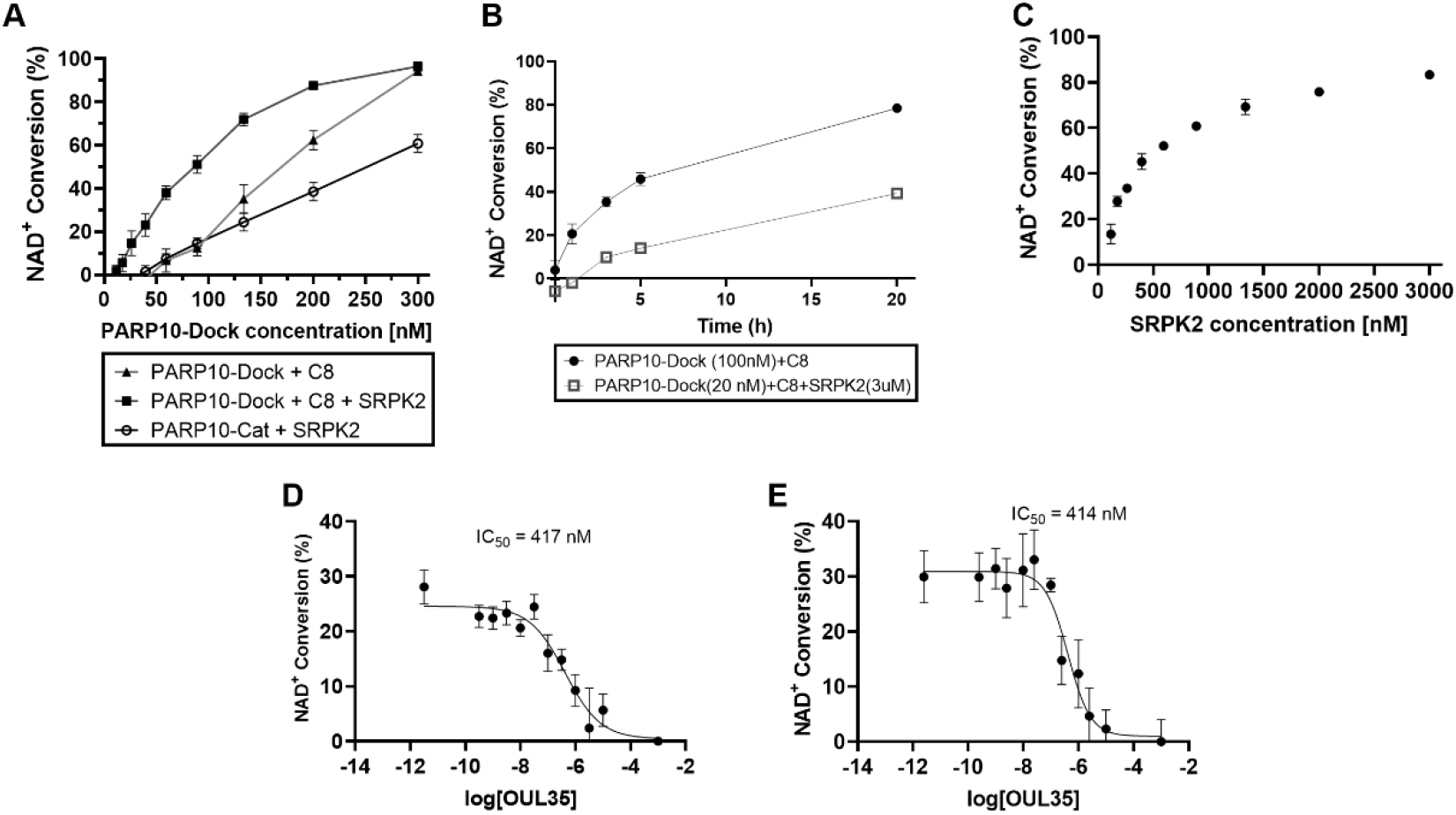
Optimization and validation of PARP10-Dock assay. **(A)** NAD^+^ conversion as a function of enzyme concentration. Assay carried out for 4h, values correspond to average ± SD of 4 repeats. **(B)** Time dependence of PAR10-Dock NAD^+^ conversion. PARP10-Dock was used at 100 nM and SRPK2 at 3 μM. The values correspond to averages ± SD **(C)** SRPK2 concentration effect on activity of 100 nM PARP10-Dock over 4 h incubation. Values correspond to conversion obtained from 3 replicates. **(D)** IC_50_ of OUL35 with PARP10-Dock+C8 using 20 nM PARP10-Dock + 3 μM SRPK2 in 50 μl wells prepared manually before 18 h incubation. **(E)** IC_50_ of OUL35 with PARP10-Dock+C8 using 150 nM enzyme prepared in 10 μl assay volume dispensed with dispensing robots. In D and E, values correspond to average ± SD of 4 replicates.

**Table I.** Conditions used for the PARPs in the proximity assay. All assays were tested using the same buffer and incubation conditions (50 mM sodium phosphate pH 7.0 during 18 h at room temperature).

|  | PARP6 | PARP7 | PARP10<br>auto | PARP10<br>manual | PARP11 | PARP12 | PARP14 | PARP15 | PARP16 |
| --- | --- | --- | --- | --- | --- | --- | --- | --- | --- |
| SRKP2 [μM] | | - | - | 2 | - | - | - | - | - |
| PARP-Dock [nM] | 80 | 300 | 120 | 20 | 80 | 80 | 140 | 200 | 3000 |
| Incubation [h] | 18 | 18 | 18 | 18 | 18 | 18 | 18 | 18 | 18 |
| Automated | Yes | Yes | Yes | No | Yes | Yes | Yes | Yes | Yes |

### Incorporation of other mono-ARTs to the scaffold assay

Finally, we extended the same scaffold incorporation strategy to other selected human mono-ARTs. For this, we generated analogous constructs containing MBP-PARP catalytic fragment-Dockerin, except for PARP14 that also included the WWE domain, which is required for robust activity (Venkannagari *et al*., 2016). As for PARP10-Dock, we studied the NAD^+^ conversion activity of each enzyme to empirically determine the optimal scaffold ratio (**Fig. S2**) and measured enzyme incorporation to scaffold by using analytical SEC (**Fig. S3**). The results showed that PARP7 was not affected by C8 presence while the optimal PARP-Dock:scaffold ratio was 0.7:1 for PARP11-Dock, 1:1 for PARP12-Dock, 1.1:1 for PARP14-Dock, 1.25:1 for PARP15-Dock and no optimal was found for PARP16 that showed no maximum.

Comparison of the NAD^+^ conversion activity of the different PARP-Cat, PARP-Dock, PARP-Dock+C8 (**Fig. 5A**) was used together with Western Blot analysis using nanoLuc-fused eAF1521 to detect MARylation (**Fig. 5B**) to assess the effect of the scaffold. PARP6 activity was increased upon complex formation with C8 scaffold (33 ± 7% vs 71 ± 7% upon scaffold addition) but SRPK2 did not promote its activity (57 ± 8%) despite being modified according to the western blot (**Fig. 5A,B**). PARP7 showed no increase in activity due to scaffold binding (51 ± 4% vs 57 ± 3%) despite it is capable of MARylate it. PARP11-Dock increased its NAD^+^ conversion activity upon interaction with C8 scaffold (from 29 ± 2% to 49 ± 3%) although the extent of scaffold modification was lower than that achieved by PARP7-Dock suggesting that the improvement in activity may originate in the proximity between the catalytic domains induced by the scaffold. Comparison to PARP7-Cat and PARP11-Cat was not performed since only they could not be produced in a soluble form.

**Fig. 5.**
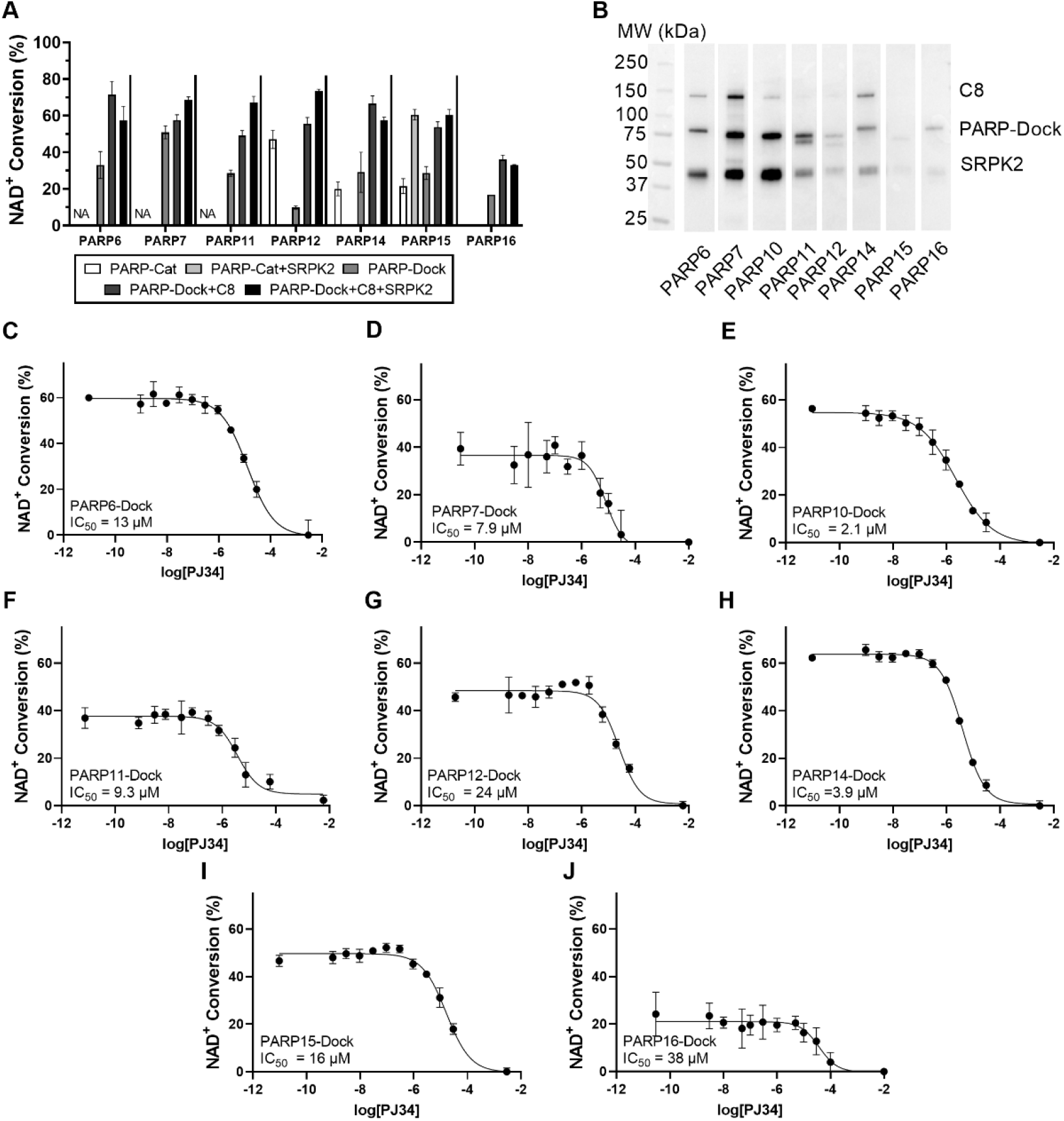
Conversion of different mono-ARTs to the scaffold-based assay. **(A)** Comparison of NAD^+^ conversion activity of different mono-ARTs when incorporated into the C8 scaffold. SRPK2 can further boost the conversion of PARP7, PARP10 and PARP11. NA indicates that the assay was not successful with the PARP-Cat construct. **(B)** Western blot analysis of components being MARylated during the assay with the different mono-ARTs. Full blots in **Fig. S4**. **C-K** IC_50_ of PJ34 inhibitor against **(C)** PARP6-Dock, **(D)** PARP7-Dock, **(E)** PARP10-Dock, **(F)** PARP11-Dock, **(G)** PARP12-Dock, **(H)** PARP14-Dock, **(I)** PARP-15-Dock and **(J)** PARP16-Dock. Values correspond to average ± SD of 4 repeats.

The PARP12-Dock construct showed clearly enhanced activity compared to PARP12-Cat (47 ± 5% with 1 μM PARP12-Cat vs. 23 ± 2% with 200 nM PARP12-Dock), and its activity was further enhanced by the addition of C8 scaffold (89 ± 4%). Altogether, the PARP12-Dock incorporated into the scaffold allows to reduce the needed protein concentration 10-fold and greatly benefitting from this approach. PARP14-Dock showed similar NAD^+^ conversion level than the PARP14-Cat (20 ± 4% and 29 ± 11%, respectively) but its activity was doubled upon incorporation in the scaffold. PARP15-Dock showed similar activity than PARP15-Cat (21 ± 4% vs. 29 ± 4%). This activity was increased upon incorporation to C8 scaffold to a similar level than the one obtained with PARP15-Cat and SRPK2 (60 ± 3% and 54 ± 3%). We speculate that this effect arises from further stabilization of PARP15-Dock upon binding to the scaffold since PARP15, in the absence of its accessory domains, primarily catalyses the hydrolysis of NAD^+^ instead of the ADP-ribose transfer to a target protein (Venkannagari *et al*., 2013). This behaviour is evidenced by the lack of strong MARylation bands in the WB which precludes the forced self-modification as the origin of the improved activity. PARP16-Dock also showed improved NAD^+^ conversion compared to PARP16-Cat (not stable activity to 17%) and its activity was slightly increased upon incorporation to the scaffold (36 ± 2%).

We finally tested whether the addition of SRPK2 to the different samples could also promote the activity of this proteins. Despite almost all the proteins could MARylate SRPK2 (**Fig. 5B**) the effect on activity varied among the proteins. A moderate increase in activity was observed for PARP7-Dock and PARP11-Dock, while it had no effect on PARP15-Dock and PARP16-Dock activity and even reduced the measured conversion of PARP12-Dock and PARP14-Dock.

Finally, we determined optimal protein concentration for the different mono-ARTs to attain 30% conversion time from time series (**Fig. S5**). The conditions for each assay were optimized also targeting the use of similar conditions across all enzymes to facilitate the profiling experiments and are summarized in **Table I**. Validatory dose response curves were measured to determine the IC_50_ of PJ34, a known inhibitor (**Fig. 5C-J**), showing in general comparable values to previously reported results (Wigle *et al*., 2020). Largest deviation from the reported values was observed for PARP16 (38 μM here vs. 0.8 μM reported).

## Discussion

In this work we used an artificial scaffold based on *H. thermocellum* cellulosome to generate a self-assembled complex containing mono-ART catalytic domains in order to improve their activity for drug discovery. We selected a cellulosome scaffold based on their properties, ease of engineering, availability and because this scaffold was shown to be easily expressed and purified using standard methods. The used scaffold incorporates flexible proline and threonine rich linkers between cohesins of 13 residues (Galera-Prat *et al*., 2018). Furthermore, the dockerin used not only provides high affinity to the scaffold cohesins but in solution is able to bind in two different orientations on the same cohesin (Vera *et al*., 2021). Altogether, these properties provide enough flexibility for binding multiple enzymes on the same scaffold and maintain them in close proximity.

The assay has been developed for PARP10 and applied to other mono-ARTs except for PARP3 and PARP4 since the original assay already works at low protein concentration. PARP9 and PARP13 were also excluded since PARP9 only modulates the activity of DTX3L (Yang *et al*., 2017; Chatrin *et al*., 2020; Ashok *et al*., 2021) and PARP13 is classified as not catalytically active (Karlberg *et al*., 2015). We could not test it with PARP8 since no soluble protein was obtained with this strategy. The use of the scaffold-based system results in an improved activity for tested mono-ARTs boosting their activity beyond 10-fold in the case of PARP12. The origin of this improved activity appears to be based on multiple factors. The fusion to MBP and dockerin modules already result in enhanced NAD^+^ conversion in many of the proteins. Since these modules do not appear to be MARylated, the most likely explanation for this observation is an increased solubility or stability of the fusion proteins compared to the isolated catalytic domains. Proximity of the catalytic units also appears to contribute to the enhanced activity of the system, as observed especially for PARP10. The bell shaped curved observed for most mono-ARTs in the NAD^+^ conversion as a function of scaffold ratio (**Fig. 3A**, **Fig. S2**) might also be interpreted in terms of enzyme proximity: high enzyme:scaffold ratio leads to most enzyme in solution, while low enzyme:scaffold ratio leads to less enzymes per scaffold and therefore each enzymatic unit is further away from others. In the case of PARP10, this leads to higher auto-modification levels (**Fig. 3D**) and the presence of its substrate protein SRPK2 also contributes to an additional increase of the NAD^+^ conversion of the system. We evaluated the possibility of forcing the proximity between PARP10 and SRPK2 by co-incorporating SRPK2-Dock in the scaffold together with PARP10-Dock to further improve the activity of the complex. Nevertheless, no additional increase was observed under these conditions (**Fig. S7**). For the other studied mono-ARTs, addition of SRPK2 showed only small beneficial to detrimental effects, despite most of the enzymes seem to be able to MARylate it to some extent, and its use was therefore not further studied.

The incorporation of additional elements in the assay system can also introduce other limitations, particularly if a candidate compound would interfere with dockerin-cohesin interaction, since this would release the enzymes from the scaffold and reduce the NAD^+^ conversion. Since dockerin requires calcium to fold and to bind cohesin, chelating compounds could potentially affect the outcome of the assay. Nevertheless, once the cohesin-dockerin interaction is established, the complex formed is very stable and, under non-denaturing conditions, can only be disassembled with high chelating agent concentrations like 25 mM EDTA at 60°C (Morag *et al*., 1996) or only partially during a gradient elution to 250 mM EDTA (Karpol *et al*., 2009). For the assay, this suggests that mixing the dockerin containing enzyme with the scaffold before adding the compound is always recommended.

The sensitivity of the homogeneous assay has limited its use to relatively high enzyme concentrations which in turn limits the possibility to measure high potency compounds. Fusion to MBP and dockerin as well as the integration of the enzymes to the scaffold to take advantage of the induced proximity leads to enhanced NAD^+^ conversion activity. The scaffold-based assay has been optimized to be performed under similar conditions for all mono-ARTs in order to facilitate profiling. The increase in activity obtained under these conditions allows to identify more potent compounds, facilitated the automation of assay dispensing thus allowing to work with large compound libraries, reducing the assay volume and lowering its cost.

## Acknowledgements

Mariano Carrión-Vázquez for constructs containing dockerin and the C1 and C8 scaffolds. The use of the facilities and expertise of the Biocenter Oulu Structural Biology core facility (a member of Biocenter Finland, Instruct-ERIC Centre Finland and FINStruct), Proteomics and Protein Analysis core facility (a member of Biocenter Finland) and Biocenter Oulu sequencing center are gratefully acknowledged.

