## Supplementary information for "Protein engineering approach to enhance activity assays of mono-ADP-ribosyltransferases through proximity"

**CONTENT**

**Table S1:** Summary of constructs

**Table S2**: Expression and purification conditions for each construct.

**Fig. S1:** Purified proteins

**Fig. S2:** Controls other PARPS (Scaffold:enzyme ratio)

**Fig. S3**: Controls other PARPS (SEC)

**Fig. S4**: Complete blots

**Fig. S5**: Other PARPs concentration dependence

**Fig. S6**: Tolerance to DMSO

**Fig. S7:** SRPK2 co-incorporation

**Table S1. Dockerin containing constructs.**

| **Construct** | **Region** | **Comments** |
|---|---|---|
| Dockerin | 671-724 | From *H. thermocellum* Cel48S |
| PARP6-Dock | 329-620 | Catalytic domain |
| PARP7-Dock | 449-655 | Catalytic domain |
| PARP10-Dock | 809-1016 | Catalytic domain |
| PARP11-Dock | 123-338 | Catalytic domain |
| PARP12-Dock | 489-701 | Catalytic domain |
| PARP14-Dock | 1535-1801 | WWE + Catalytic domain |
| PARP15-Dock | 481-678 | Catalytic domain |
| PARP16-Dock | 1-279 | Catalytic domain |
| SRPK2-Dock | 51-688 | |
| C1 | 29-182 | From *H. thermocellum* CipA cohesin 1 + linker to cohesin2 |
| C8 | 29-182 | 8 tandem repeats separated by sequence AS |

**Table S2. Expression and purification conditions.**

| Construct | Expression strain | Media | Expression Conditions | Purification steps |
|---|---|---|---|---|
| PARP6-Dock | Rosetta 2 | TB-AIM + 0.8% glycerol + 10 mM Benzamide | 16°C, 25 h, 2 mM $CaCl_2$ | IMAC (5 ml Column) + MBP-trap + SEC |
| PARP7-Dock | Rosetta 2 | TB + 0.8% glycerol + 10 mM Benzamide | 18°C, 16 h, 0.2 mM IPTG, 2 mM $CaCl_2$ | IMAC (resin) + SEC |
| PARP10-Dock | BL21 | TB-AIM + 0.8% glycerol | 18°C, 16 h, 2 mM $CaCl_2$ | IMAC (5 ml Column) + SEC |
| PARP11-Dock | Rosetta 2 | TB + 0.8% glycerol | 37°C, 3 h, 0.15 mM IPTG, 2 mM $CaCl_2$ | IMAC (resin) + SEC |
| PARP12-Dock | BL21 | TB-AIM + 0.8% glycerol | 18°C, 16 h, 2 mM $CaCl_2$ | IMAC (5 ml Column) + SEC |
| PARP14-Dock | BL21 | TB-AIM + 0.8% glycerol | 18°C, 16 h, 2 mM $CaCl_2$ | IMAC (resin) + SEC |
| PARP15-Dock | BL21 | TB-AIM + 0.8% glycerol | 18°C, 16 h, 2 mM $CaCl_2$ | IMAC (resin) + SEC |
| PARP16-Dock | BL21 | TB-AIM + 0.8% glycerol | 18°C, 16 h, 2 mM $CaCl_2$ | IMAC (5 ml Column) + MBP + SEC |
| SRPK2-Dock | BL21 | TB-AIM + 0.8% glycerol | 18°C, 16 h, 2 mM $CaCl_2$ | IMAC (5 ml Column) + SEC |
| C1 | BL21 | TB-AIM + 0.8% glycerol | 14°C, 20 h | Heat incubation + IMAC (5 ml Column) |
| C8 | C41 | TB-AIM + 0.8% glycerol | 14°C, 20 h | Heat incubation + IMAC (5 ml Column) |

| | | | | |
|---|---|---|---|---|
| PARP10-Cat | BL21 | TB-AIM + 0.8% glycerol | 18°C, 16 h | IMAC (5 ml Column) + SEC (Venkannagari *et al.*, 2013) |
| PARP12-Cat | BL21 | TB-AIM + 0.8% glycerol | 18°C, 16 h | IMAC (5 ml Column) + SEC (Venkannagari *et al.*, 2016) |
| PARP14-Cat | Rosetta 2 | TB-AIM + 0.8% glycerol | 18°C, 16h | IMAC (2x1ml Column) + SEC (Venkannagari *et al.*, 2016) |
| PARP15-Cat | Rosetta 2 | TB-AIM + 0.8% glycerol | 18°C, 16h | IMAC (1 ml Column) + SEC (Venkannagari *et al.*, 2013) |
| PARP16-Cat | BL21 | TB-AIM + 0.8% glycerol | 18°C, 16 h | IMAC (5 ml Column) + SEC (Venkannagari *et al.*, 2016) |
| SRPK2 | BL21 | TB-AIM + 0.8% glycerol | 18°C, 16 h | IMAC (5 ml Column) + SEC (Venkannagari *et al.*, 2016) |
| Dockerin | BL21 | TB-AIM + 0.8% glycerol | 18°C, 16 h | IMAC (5 ml Column) + SEC + TEV cleavage + reverse IMAC |
| MBP | BL21 | TB-AIM + 0.8% glycerol | 18°C, 16 h | IMAC (5 ml Column) + SEC + TEV cleavage + reverse IMAC |
| nLuc-eAF1521 | BL21 | TB-AIM + 0.8% glycerol | 18°C, 16 h | IMAC (5 ml column)(Sowa *et al.*, 2021) |

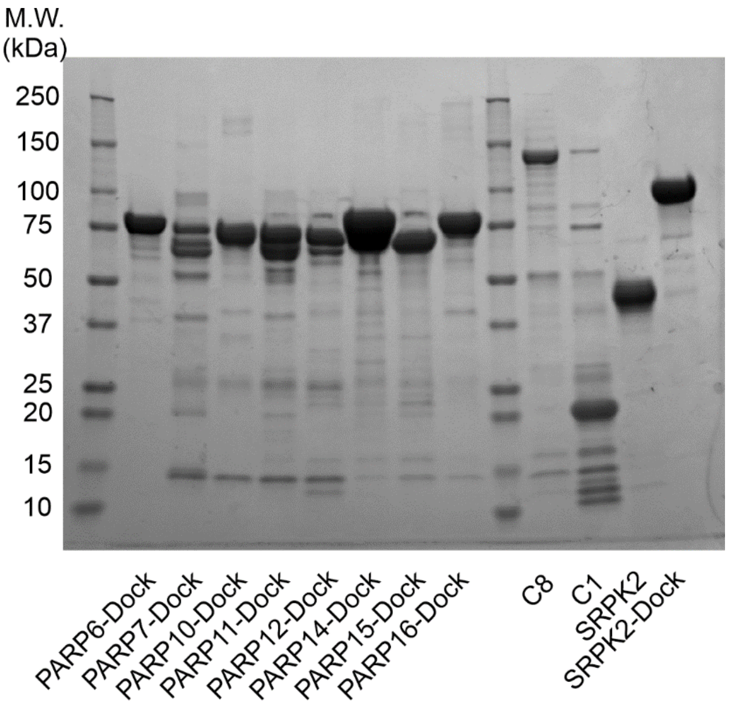


**Fig. S1:** SDS- PAGE of purified proteins used in the scaffold-based assay.

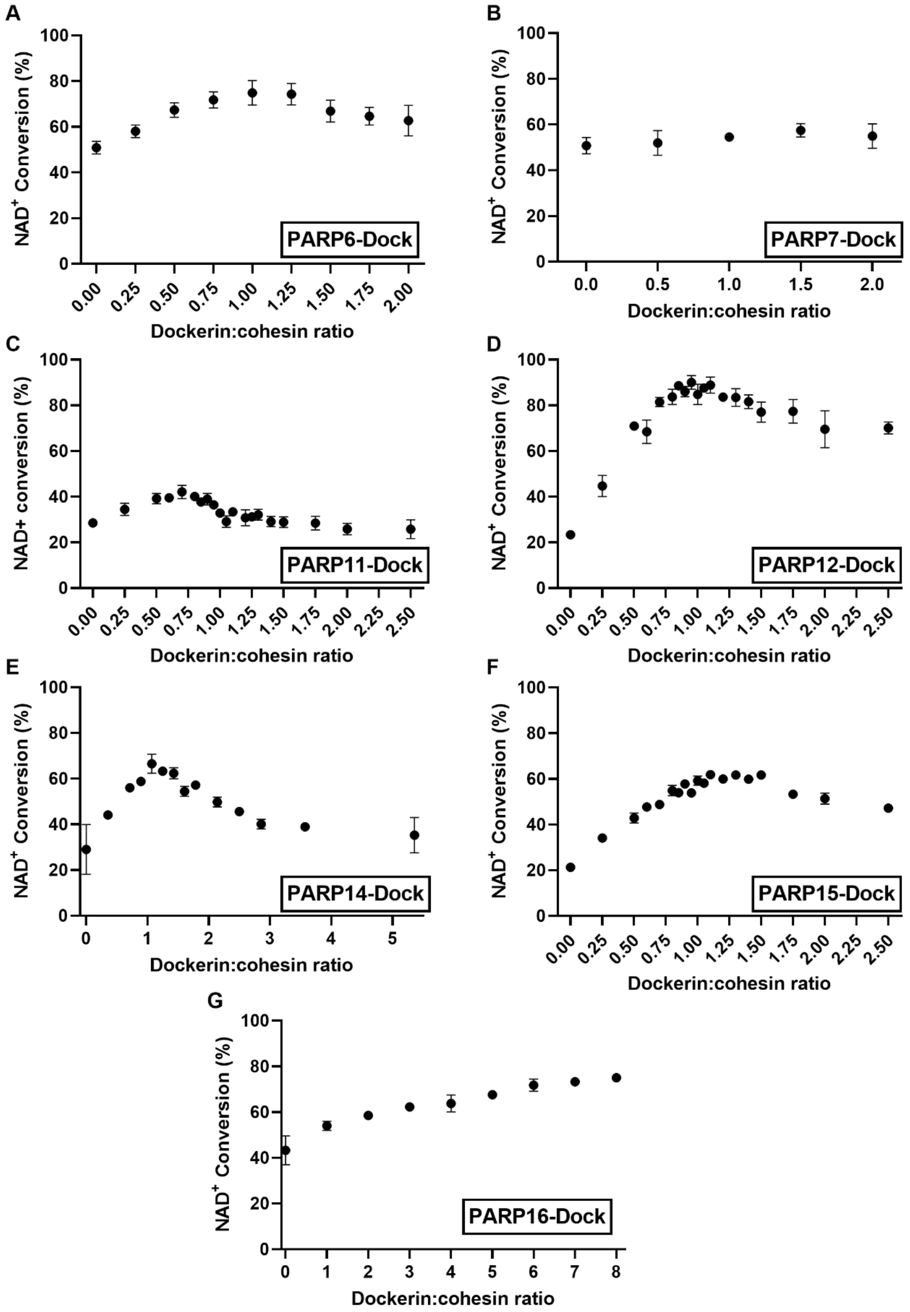


**Fig. S2:** Determination of PARP-Dock:scaffold optimal ratio for different PARP-Dock. **(A)** PARP6-Dock at 120 nM **(B)** PARP7-Dock at 500 nM. **(C)** PARP11-Dock at 80 nM, **(D)** PARP12-Dock at 200 nM. **(E)** PARP14-Dock at 140 nM. **(F)** PARP15-Dock at 200 nM. **(G)** PARP16-Dock at 1.6 µM. The optimal scaffold:enzyme concentration was determined from the maximum conversion conditions. That ratio was used in all other experiments with each PARP-Dock. Values represent average ± SD of 4 replicates.

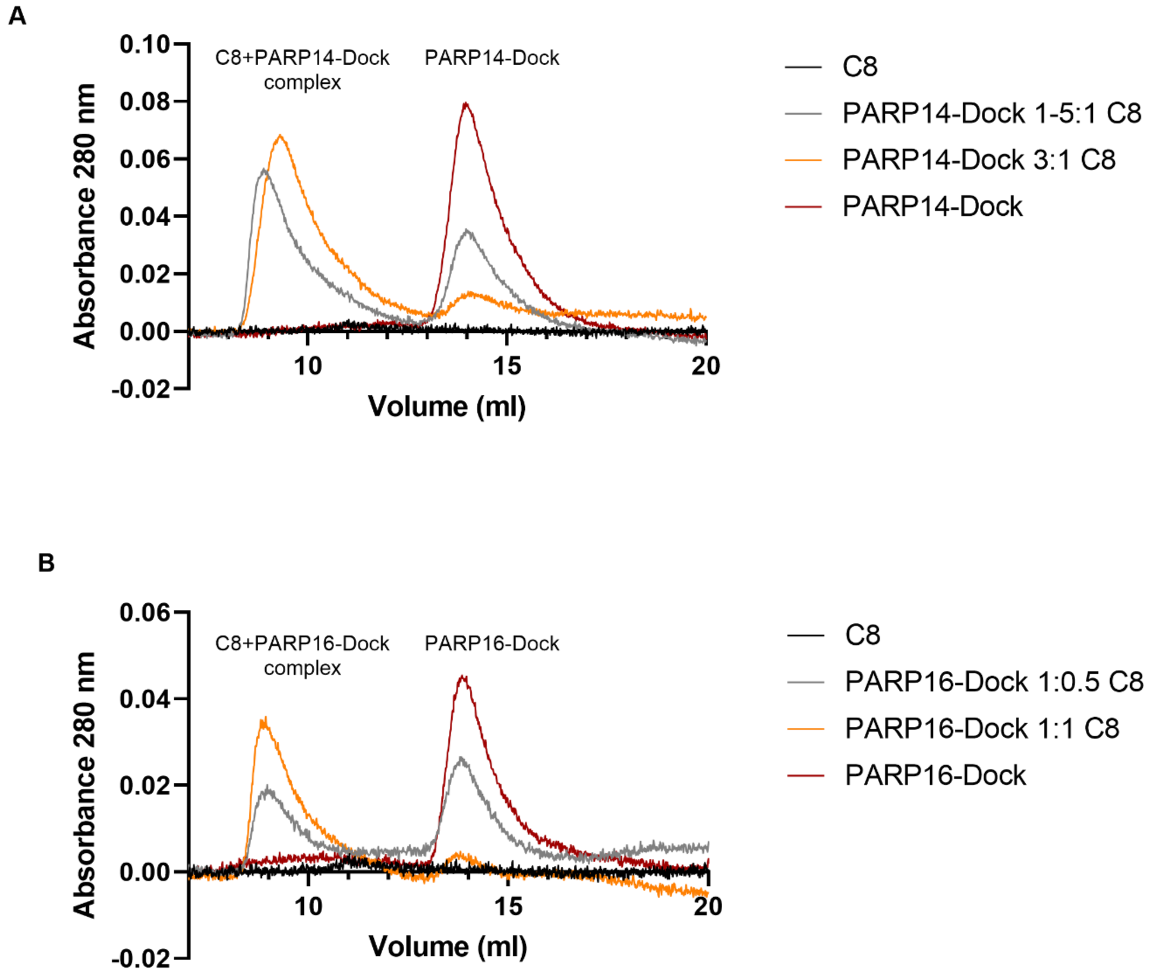


**Fig. S3**: Complex formation studied by SEC at the ratios where activity is optimal. **(A)** PARP14-Dock can form a complex with C8 scaffold (yellow). When the scaffold concentration is halved, excess unbound PARP14-Dock can be observed (gray). **(B)** PARP16-Dock can be incorporated in the C8 scaffold (yellow). When the scaffold concentration is halved, excess unbound PARP16-Dock can be observed (gray).

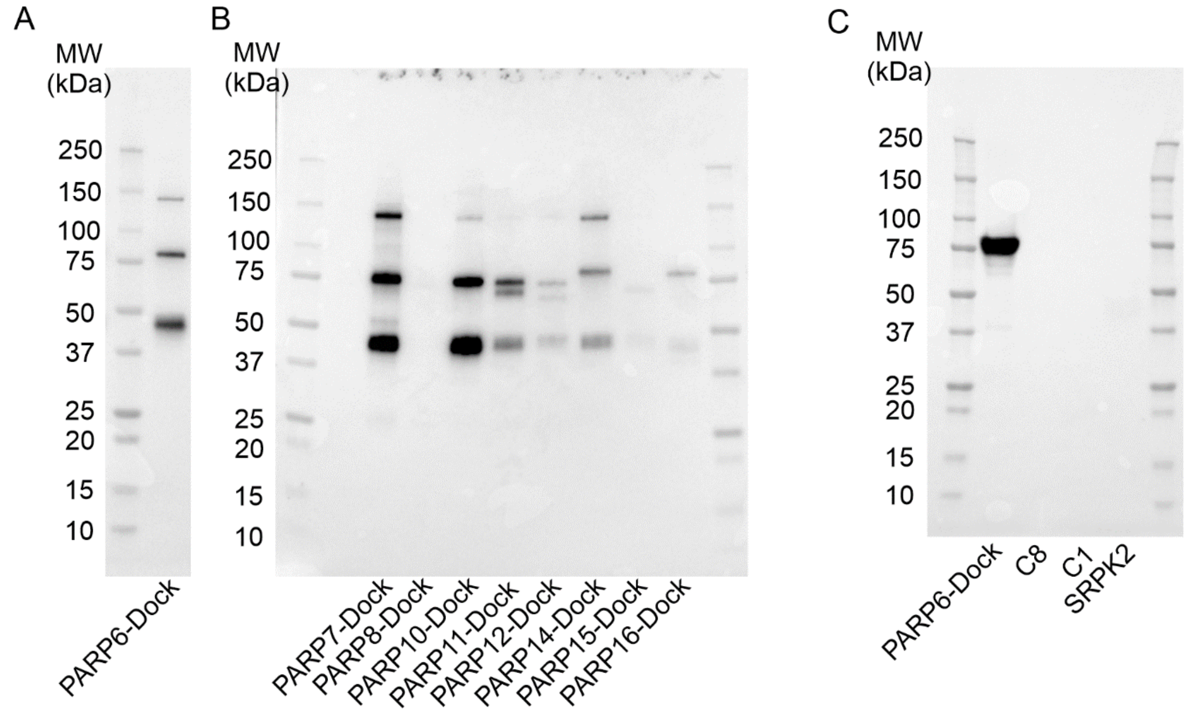


**Fig. S4:** Full blots from fragments displayed in figure 5. **(A)** Western blot analysis of PARP6-Dock incubated with SRPK2, C8 scaffold and $NAD^+$. **(B)** Full western blot of the other PARP-Dock incubated with SRPK2, C8 scaffold and $NAD^+$. PARP8-Dock consist of region 611-844 of PARP8, this construct was mostly insoluble and not active. **(C)** Control western blot where PARP6-Dock, C8, C1 or SRPK2 were incubated in the presence of $NAD^+$. Western blots were proved with nanoLuc-eAF1521(Sowa *et al.*, 2021).

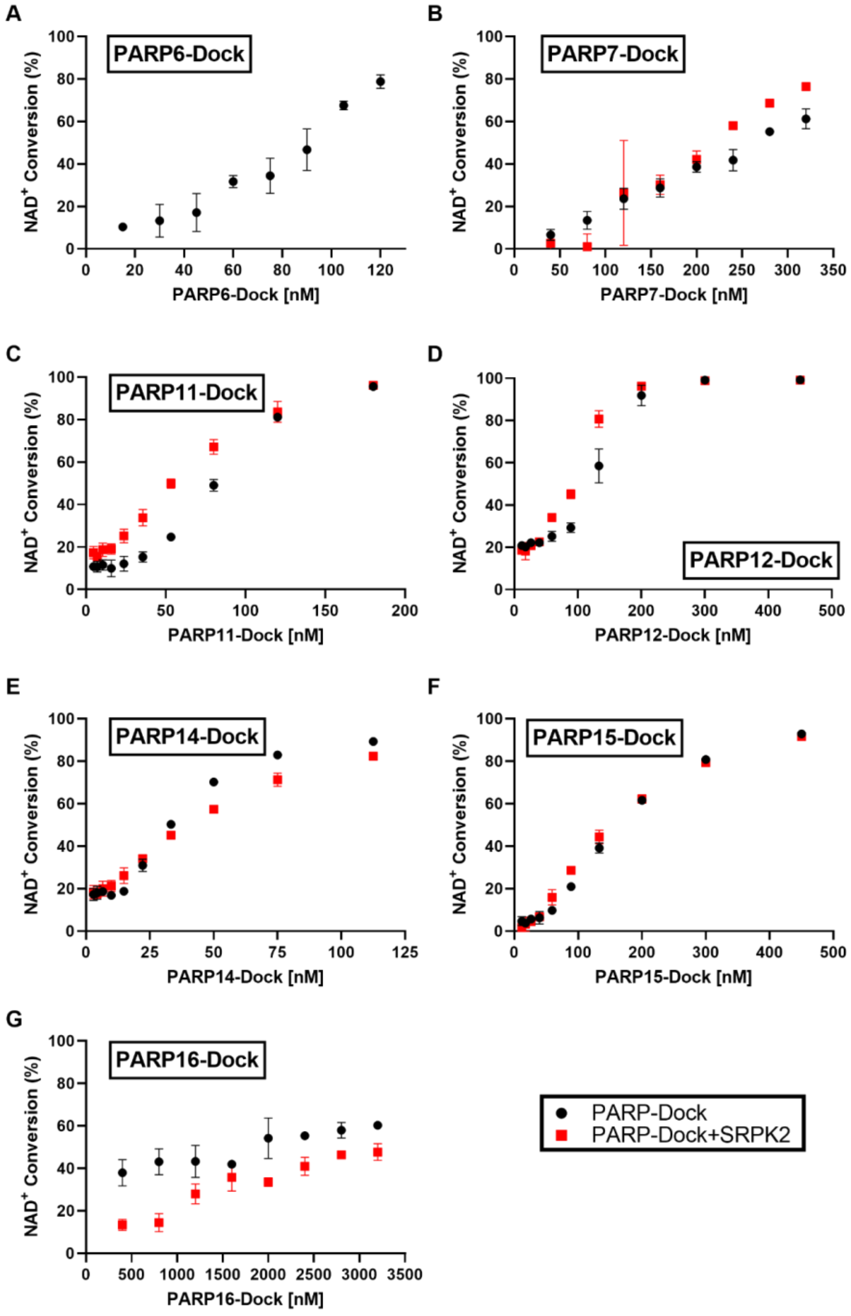


**Fig. S5**: PARP-Dock NAD$^+$ conversion activity dependence on enzyme concentration. Converted NAD$^+$ for **(A)** PARP6-Dock, **(B)** PARP7-Dock, **(C)** PARP11-Dock, **(D)** PARP12-Dock, **(E)** PARP14-Dock, **(F)** PARP15-Dock and **(G)** PARP16-Dock in the presence of C8 scaffold. SRPK2 was used at 500 nM. Concentration to reach 30% conversion was used in subsequent experiments. Reactions were carried out for 19-20 h at RT and values represent average ± SD of 4 replicates, except for PARP16-Dock where incubation time was 3 h and 2 replicates where measured.

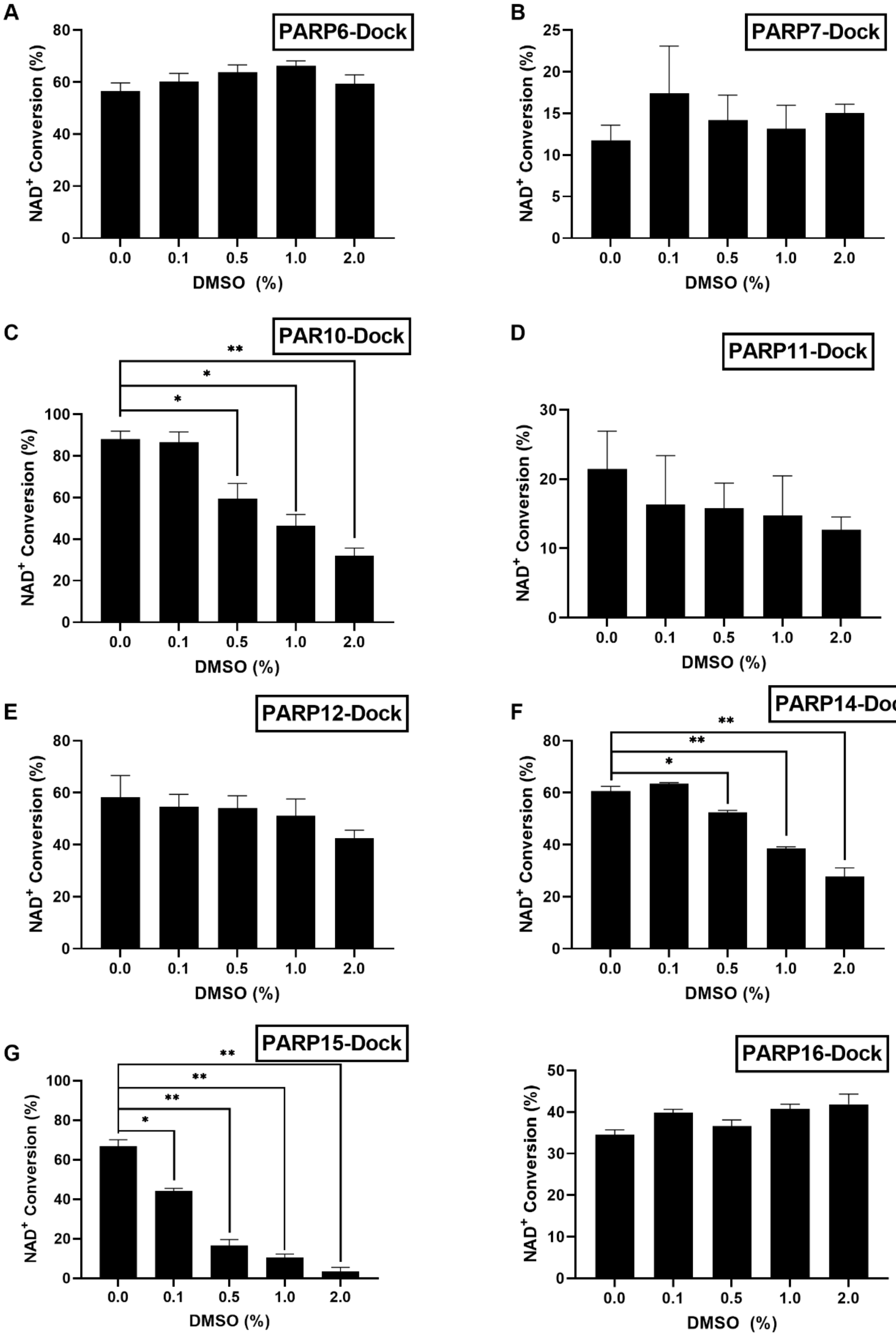


**Fig. S6**. Tolerance of mono-ART-Dock $NAD^+$ conversion activity to DMSO presence. **(A)** 100 nM PARP6-Dock, **(B)** 200 nM PARP7-Dock, **(B)** 150 nM PARP10-Dock, **(C)** 60 nM PARP11-Dock, **(D)** 150 nM PARP12-Dock, **(E)** 30 nM PARP14-Dock, **(F)** 80 nM PARP15-Dock, **(G)** 3 µM PARP16-Dock. Values are presented as average ± SD of duplicates.

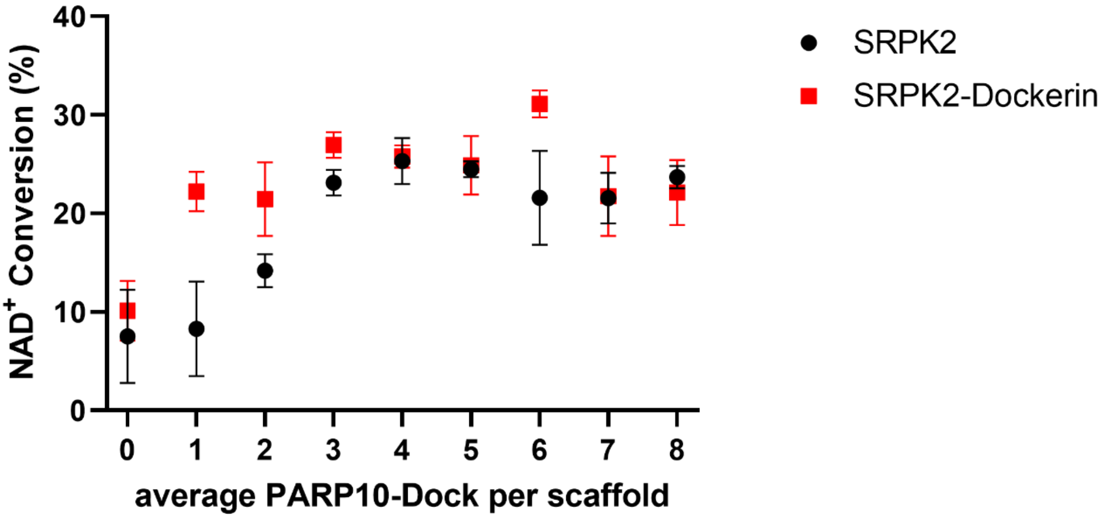


**Fig. S7:** SRPK2-Dock co-incorporation on scaffold together with PARP10-Dock does not increase PARP10-Dock $NAD^+$ conversion. 50 nM PARP10-Dock was incubated with 500 nM SRPK2 or SRPK2-Dock and varying ratios of C8 scaffold to produce complexes with different average number of enzymatic units per scaffold. Empty scaffold places can be occupied by SRPK2-Dock or kept empty when SRPK2 is used. Values correspond to the average ± SD of quadruplicates.

The use of complexes with average low PARP10-Dock per scaffold results in a decrease of the activity of the system when SRPK2 is present, consistent with a decreased proximity of enzymes. When SRPK2-Dock is used, the activity is maintained to the level of the fully saturated scaffold even when a single PARP10-Dock is present per each scaffold molecule suggesting that SRPK2-Dock in the scaffold is preferentially MARylated. In the absence of scaffold, both SRPK2 and SRPK2 conditions show similar conversion.
